# Genomic islands of divergence between *Drosophila yakuba* subspecies predominantly overlap with chromosomal inversions

**DOI:** 10.1101/2022.07.16.500113

**Authors:** Erina A. Ferreira, Cathy C. Moore, David Ogereau, Arnaud Suwalski, Stéphane R. Prigent, Rebekah L. Rogers, Amir Yassin

## Abstract

During the early stages of local adaptation and speciation, genetic differences tend to accumulate at certain regions of the genome leading to the formation of genomic islands of divergence (GIDs). This pattern may be due to selection and/or difference in the rate of recombination. Here, we investigate the possible causes of GIDs in *Drosophila yakuba mayottensis*, and reconfirm using field collection its association with toxic noni (*Morinda citrifolia*) fruits on the Mayotte island. Population genomics revealed lack of genetic structure on the island and identified 20 GIDs distinguishing *D. y. mayottensis* from generalist mainland populations of *D. y. yakuba*. The GIDs were enriched with gene families involved in the metabolism of lipids, sugars, peptides and xenobiotics, suggesting a role in host shift. We assembled a new genome for *D. y. mayottensis* and identified five novel chromosomal inversions. Twelve GIDs (∼72% of outlier windows) fell close to or within subspecies-specific inversions. However, three GIDs were in collinear, high recombining regions indicating strong signal of hard selective sweeps. Unlike *D. y. mayottensis*, *D. sechellia*, the only other noni-specialist, is homosequential with its generalist relatives. Thus, both selection and rearrangements shape GIDs and striking convergences can occur between species with distinct genomic architectures.

## Introduction

The genomic landscape of inter-population differentiation is often heterogeneous, with certain genomic regions, known as genomic islands of divergence (GIDs), emerging above the average differentiation level (Feder and Nosil 2010; Wolf and Ellegren 2017). GIDs were originally hailed as a result of divergent selection acting on genes that underlie adaptation to distinct environment and/or reproductive isolation (Turner et al. 2005). However, a plethora of mechanisms may also lead to the formation of such islands even in the absence of divergent selection. Most importantly, variation in recombination rate, due to heterochromatin content or structural variation, strongly affects the rate of admixture and the accumulation of genetic differences in natural populations (Feder and Nosil 2010; Cruickshank and Hahn 2014; Quilodrán et al. 2020). Chromosomal inversions are among the most common forms of structural variation, and a number of studies has unraveled an overlap between inversions and GIDs (Feder and Nosil 2009; Michel et al. 2010; Ellegren et al. 2012; Sodeland et al. 2016; Huang et al. 2020). Inversions often suppress recombination, reducing the rate at which slightly-deleterious mutations are purged from the population (Berdan et al. 2021; Jay et al. 2021, 2022). Conversely, the reduction of recombination could conserve one or more favorable allelic combinations for survival in contrasting environments, hence becoming targets of balancing or diverging selection depending on the heterogeneity of the environment (Faria et al. 2019; Kapun and Flatt 2019; Mérot et al. 2021; Hager et al. 2022; Harringmeyer and Hoekstra 2022; Sanchez-Donoso et al. 2022; Tepolt et al. 2022). Untangling the complex interactions of selection and inversions in driving GIDs and ultimately speciation therefore requires proper understanding of the genomic architecture of the study model and its natural variation.

Host shifts or changes in the width of the ecological niche, e.g., specialization, are a major source of diversification in herbivorous insects (Wiens et al. 2015; Forbes et al. 2017). The complexity of this phenomenon, which should involve the simultaneous evolution of genes underlying multiple behavioral and physiological phenotypes with an ultimate impact on reproductive isolation, has long been a major area of research (Futuyma and Moreno 1988; Vertacnik and Linnen 2017). One of the best well-characterized systems at the genomic level is the specialization of the fly *Drosophila sechellia* on toxic fruits of noni (*Morinda citrifolia*) on the Seychelles islands in the Indian Ocean (Louis and David 1986; Auer et al. 2021). *Drosophila sechellia* is a member of the *simulans* species complex, which comprises two additional species, namely the cosmopolitan *Drosophila simulans* and *Drosophila mauritiana*, an endemic to Mauritius. Unlike *D. sechellia*, both species are generalists and are repulsed by the odor of and intolerant to noni main toxins, octanoic and hexanoic acids (R’Kha et al. 1991; Amlou et al. 1997, 1998; Drum et al. 2022). The three species represent a sister clade to *Drosophila melanogaster*. This facilitated the extension of the unique genetic and genomic tools from this paradigmatic species to the study of *D. sechellia* specialization (Matsuo et al. 2007; Lavista-Llanos et al. 2014; Prieto-Godino et al. 2017; Auer et al. 2020). Besides, the three species show partial reproductive isolation and are karyologically homosequential, making them ideal for quantitative trait loci (QTLs) mapping through interspecific crosses (Earley and Jones 2011; Hungate et al. 2013; Huang and Erezyilmaz 2015; Chakraborty et al. 2021). Despite these numerous advantages, the divergence of *D. sechellia* from its generalist sister species *D. simulans* is estimated at around 89.000-242,000 years ago (Garrigan et al. 2012; Schrider et al. 2018); its entire genome has accumulated so many differences that traces of possible GIDs during the early stages of divergence may be difficult to observe. Besides, a particularly low level of genetic diversity has been detected in this species (Legrand et al. 2009; Schrider et al. 2018), further complicating population genomics analyses.

In January 2013, a second case of *Drosophila* species associated with toxic noni was discovered in Mayotte, another island in the Indian Ocean. This was a population of *Drosophila yakuba*, which had higher preference for and survival on noni unlike its continental conspecifics (Yassin et al. 2016). The noni-specialist population likely diverged from generalist continental populations nearly 28,000 years ago and specialization was accompanied by precopulatory isolation between Mayotte and continental flies, earning them subspecific status as *D. y. mayottensis* and *D. y. yakuba*, respectively (Yassin et al. 2016; Yassin 2017). Genomic comparisons between this single population of *D. y. mayottensis* and two *D. y. yakuba* continental populations identified multiple GIDs, widespread across the genome, consistent with theoretical expectations (Yassin et al. 2016). Many *D. y. mayottensis* GIDs overlap with large noni survival QTLs in *D. sechellia* (Yassin et al. 2016). Despite those similarities, *D. yakuba* is a species where, unlike the *simulans* species complex, multiple polymorphic chromosomal inversions have been identified in continental populations (Lemeunier and Ashburner 1976). Several GIDs in *D. y. mayottensis* may therefore be the result of such inversions.

In this paper, we report the population genomics analysis of three additional populations of *D. y. mayottensis* all associated with noni in Mayotte. We identified 20 GIDs distinguishing the three populations from continental *D. y. yakuba*. We assembled a genome for *D. y. mayottensis* and identified multiple inversions that predominantly overlap with the GIDs, hence allowing the discrimination between GIDs presenting genuine signal of hard selective sweep from those which may be mere results of structural variation.

## Materials and methods

### Fly collection and establishment of isofemale lines from Mayotte island

Flies were prospected in April 2017 from 14 localities on the Grande Terre and Petite Terre islands of Mayotte. We particularly searched fruiting noni trees but also collected drosophilids in places with no noni trees. Flies were collected by net sweeping and direct aspiration over fallen fruits or using standard baits with fermented banana hung from tree branches. Flies were also bred from ripe fruits brought back to the laboratory. *D. y. mayottensis* isofemale lines were established by placing one female and one male in a small vial with instant Carolina medium and the F_1_ progeny was transferred to an axenic medium in the laboratory. Fly species that were or could not be bred in the laboratory were killed in absolute ethanol and taxonomically sorted in the laboratory. Data on *D. yakuba* seasonal abundance and attraction to banana baits in West Africa was obtained from Vouidibio (1985).

### Pooled and individual lines genome sequencing

*Drosophila yakuba mayottensis* were sequenced both as pooled (Pool-Seq) as well as individual lines (Ind-Seq). For Pool-Seq analyses, we pooled single F_1_ female per line in three pools according to the populations, namely BE, MTS and SOU. Pools were sent in ethanol to End2end Genomics LLC, Davis, California where DNA was extracted, PE library prepared using NEBNext Protocol and 75x coverage double-paired Illumina NovoSeq sequencing was conducted. For Ind-Seq analyses, DNA was extracted from single flies flash frozen in liquid nitrogen following QIAamp Mini Kit (Qiagen) protocol without using RNase A. The resulting DNA samples were quantified (Qubit dsDNA HS assay kit, ThermoFisher Scientific), assessed for quality (Nanodrop ND-2000), and stored at −20°C. Illumina TruSeq Nano DNA libraries were prepared manually following the manufacturer’s protocol (TruSeq Nano DNA, RevD; Illumina). Briefly, samples were normalized to 100ng DNA and sheared by sonication with Covaris M220 (microTUBE 50; Sage Science). The samples were end repaired, purified with Ampure XP beads (Agencourt; Beckman Coulter), adaptors adenylated, and Unique Dual Indices ligated. Adaptor enrichment was performed using eight cycles of PCR. Following Ampure XP bead cleanup, fragment sizes for all libraries were measured using Agilent Bioanalyzer 2100 (HS DNA Assay; Applied Biosystems). The libraries were diluted 1:10 000 and 1:20 000 and quantified in triplicate using the KAPA Library Quantification Kit (Kapa Biosystems). Equimolar samples were pooled, and the libraries were size selected targeting 400-700bp range to remove adaptor monomers and dimers using Pippen Prep DNA Size Selection system (1.5% Agarose Gel Cassette #CDF1510; Sage Sciences). Library pools were run on an Illumina HiSeq 4000 platform using the 150bp paired end (PE) Cluster Kit.

### Population genomics analyses and delimitation of the genomic islands of divergence (GIDs) in *Drosophila yakuba*

As in Yassin et al. (2016) and Ferreira et al. (2021), we used previously published genomic sequences from two continental *D. y. yakuba* populations, namely Cameroon (CY) and Kenya (NY) (Rogers et al. 2014), for comparisons with *D. y. mayottensis*. We also used previous Pool-Seq reads from a single *D. y. mayottensis* population collected from Soulou in 2013 for temporal comparisons within the subspecies (Yassin et al. 2016). For all comparisons, reads were mapped to the *D. yakuba* v.1.05 reference genome obtained from Flybase (https://flybase.org/, Thurmond et al. 2019) using Minimap2 software package (Li 2018). Minimap2-generated SAM files were converted to BAM format using samtools 1.9 software (Li et al. 2009). The BAM files were then cleaned and sorted using Picard v.2.0.1 (http://broadinstitute.github.io/picard/). Popoolation 2 software package v.1.20162 (Kofler et al. 2011) was used to generate a synchronized mpileup file, from which intra-population polymorphism (*π)* and between-populations genetic differentiation (*F_ST_*) were estimated using customized perl scripts.

### Delimitation of the genomic islands of divergence (GIDs) in *D. y. mayottensis*

We conducted Population Branch Excess (*PBE*) analysis (Yassin et al. 2016) to identify GIDs specifically distinguishing *D. y. mayottensis* from the two continental populations CY and NY. *PBE* is a standardized version of the Population Branch Statistic (*PBS*), which was proposed to identify population-specific allelic changes from *F_ST_* pairwise comparisons between three populations (Yi et al. 2010). The standardization measures how *PBS* at a locus deviates from the chromosome-wide median *PBS* divided by the median *F_ST_* estimate from the two non-focal populations. Therefore, *PBE* is less sensitive than *PBS* to genome-wide variation in polymorphism levels. Due to the lack of genetic structure within *D. y. mayottensis* (see below), Pool-Seq reads of the three populations were pooled into a single Pool of Pools (PoP). *PBS* and *PBE* were made assuming an (CY,(NY,PoP)) tree. Genes with *PBE* values falling ≥ 97.5 quantiles were considered outliers, and the width of a genomic island of divergence was considered if two outlier windows did not fall more than 500-kb apart. Gene enrichment tests were conducted for each GID as well as using the whole list of genes from all GIDs using the online Database for Annotation, Visualization and Integrated Discovery (DAVID) (Sherman et al. 2022).

### De novo assembly of *D. y. mayottensis* genome and identification of chromosomal rearrangements

A mass culture of *D. y. mayottensis* was used to assemble a reference genome for this subspecies. Genomic DNA was extracted using the Nucleobond AXG20 kit and buffer set IV from Macherey-Nagel (ref. 740544 and 740604, https://www.mn-net.com, Düren, Germany), and size selection was conducted using the SRE XS kit from Circulomics (https://www.circulomics.com/, Baltimore, Maryland, USA). Fragments less than 10 Kb were eliminated. The SQK-LSK109 kit from Oxford nanopore technology (Lu et al. 2016), https://nanoporetech.com/) was then used to prepare the samples for nanopore sequencing following manufacturer’s protocol. The library was then loaded on a R9.4 flow cell for sequencing on MinION. Raw data were basecalled with Guppy v4.0.11.

Illumina short-read sequences have also been produced to reconstruct our genome, using a hybrid assembly strategy, combining those short accurate Illumina reads to our long Nanopore reads. Illumina sequencing was realized by Novogene Company Limited (https://en.novogene.com, Cambridge, UK) on our previously extracted DNA. DNA sequences, from Nanopore and Illumina technologies, were then assembled using MaSuRCA genome assembler software v3.4.0 (Zimin et al. 2013). We aligned our newly constructed *D. y. mayottensis* genome using minimap2 software package to the v.1.05 reference genome as well as to a newly assembled genome of the reference *D. yakuba* strain (David et al. 2022) to correct for early missassemblies in the v.1.05 reference genome (Miller et al. 2018). *D. y. mayottensis* scaffolds were then ordered and oriented according to the reference genome, and chromosomal rearrangements were visually detected from a dotplot constructed by R. We used the cytological position of *D. melanogaster* orthologs of *D. yakuba* genes to delimit *D. y. yakuba* rearrangements described by Lemeunier and Ashburner (1976).

## Results

### *Drosophila yakuba mayottensis* is strictly associated with noni in Mayotte

In April 2017, we investigated the association between *D. y. mayottensis* and noni across the two main islands of Mayotte, *i.e* Grande Terre and Petite Terre. Out of 14 collection sites, we identified noni trees in 9 locations, mostly near the coastline. However, the trees were not always at the same fruiting stage, with fallen ripe fruits found at only three locations, namely Soulou (the same locality for the 2013 sample, hereafter SOU), M’tsangamouji (MTS) in the humid northwest and Bambo Est (BE) in the arid southeast (Figure 1A). *D. y. mayottensis* flies were collected through net sweeping or direct aspiration over fallen, rotting noni fruits only at those three locations. No single *D. y. mayottensis* fly was collected using fermenting banana traps at those sites or at any other site on the island, although the noni trees at Bambo Est were present amid a banana plantation (Figure 1B). In West Africa, such traps often collect tens or hundreds of *D. y. yakuba* (Rio et al. 1983; Vouidibio 1985; Llopart et al. 2005; Prigent et al. 2013; Turissini and Matute 2017) (Figure 1B; *χ*^2^ *P* < 0.001 for comparison with data from Vouidibio (1985)). The total number of collected *D. y. mayottensis* was low, ∼200 flies throughout the 10-days duration of the expedition, compared to ∼600 flies that were collected during the last 2 days of the January 2013 at a single location (David et al. 2014). This indicates a strong seasonality in the abundance of the species, consistent with other observations on *D. y. yakuba* on the continent (Vouidibio 1985; Prigent et al. 2013), and highlights the necessity of year-round investigations. We also observed that flies were more abundant on noni fruits that have been nibbled and thrown to the ground by the brown lemur (*Eulemur fulvus*) than on intact fallen fruits (Figure 1C).

**Figure 1 –.**
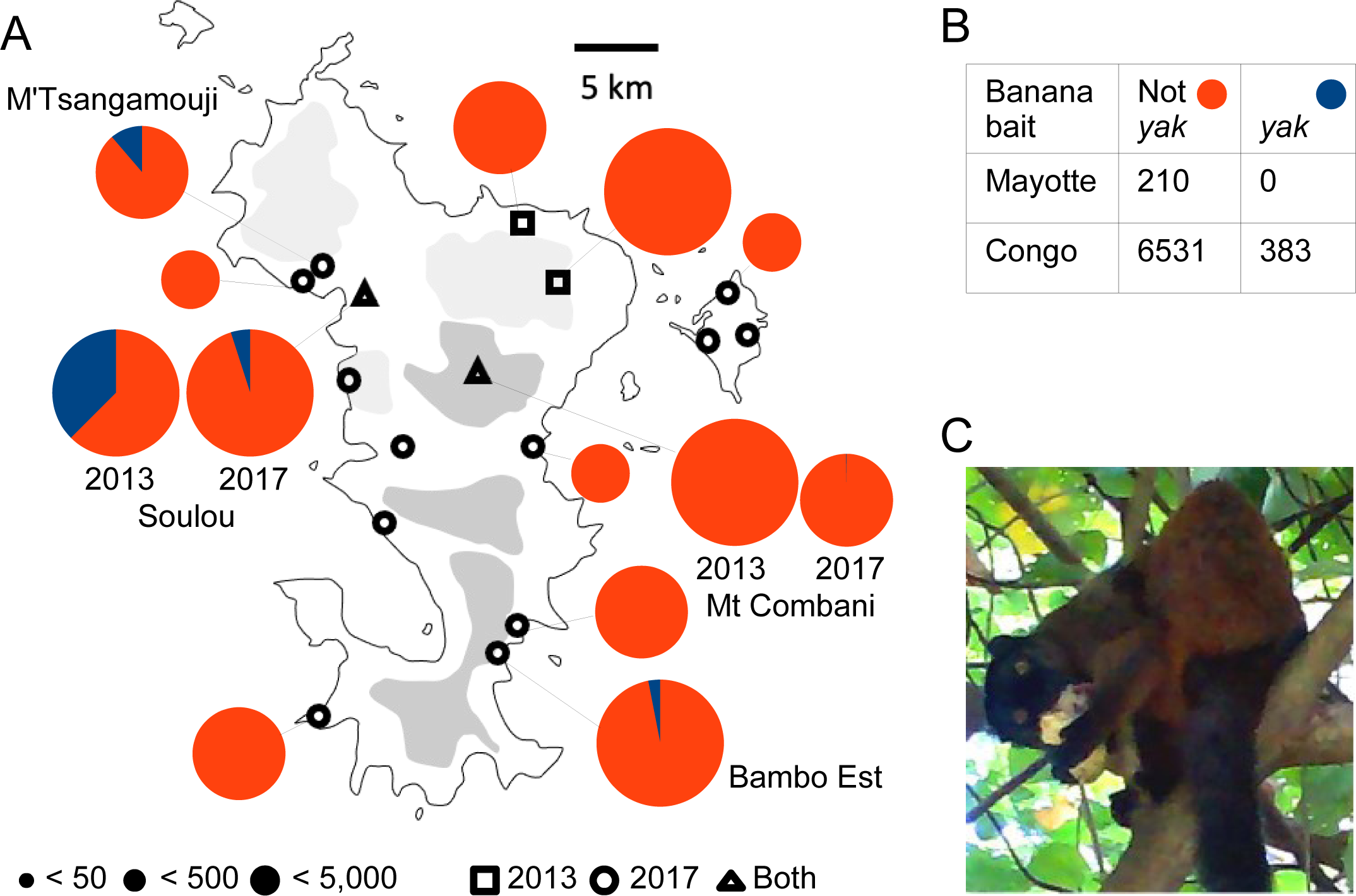
Association of *Drosophila yakuba mayottensis* with noni (*Morinda citrifolia*) on the island of Mayotte. A) Frequency of *D. yakuba* (blue) among drosophilids (red) collected on the island of Mayotte in January 2013 (after David et al. (2014)) and April 2017. Locations with fruiting noni trees are indicated with a star. The size of the pie charts reflects the total number of collected flies per site. B) Abundance of *D. yakuba* flies among drosophilids collected using standard fermenting banana traps in Congo (after Vouidibio (1985) and in Mayotte. C) A brown lemur (*Eulemur fulvus*) eating a noni fruit in Mayotte (© Amir Yassin).

### Pooled and isofemale line sequencing of the three Mayotte populations reveal low level of differentiation on the island

We established 26, 22 and 29 isofemale lines from wild-caught flies from the three collection sites, *i.e.,* BE, SOU and MTS, respectively. As in Yassin et al. (2016), we sequenced a single F_1_ fly per line in three pools according to the populations. The level of genetic differentiation estimated in 10-kb windows in SOU between the 2013 and 2017 samples was low (*F_ST_* = 0.0262 ± 0.0001; Figure 2A), as well as the level of differentiation between the three 2017 geographical samples (*F_ST_* = 0.0283 ± 0.0001; Figure 2B; Supplementary Table S1). Because *F_ST_* estimates from Pool-Seq data could be slightly biased (Hivert et al. 2018), we also sequenced 7, 6 and 6 individual isofemale lines from BE, SOU and MTS, respectively. We found a strong correspondence between *F_ST_* estimates from both approaches, with Ind-Seq data producing higher estimates probably due to their smaller sample size and higher ascertainment bias (*F_ST_* = 0.0352 ± 0.0001 between the 2013 and 2017 of SOU, and 0.0426 ± 0.0001 between the three Mayotte populations; Supplementary Table S1). Nonetheless, whether Pool- or Ind-Seq analyses were conducted, the genetic differentiation among *D. y. mayottensis* populations was significantly lower than that between the two *D. y. yakuba* populations from Cameroon (CY) and Kenya (NY) (*F_ST_* = 0.0671±0.0002; Student’s *t*-test *P* < 2.2 x 10^-16^ for both analyses).

**Figure 2 –.**
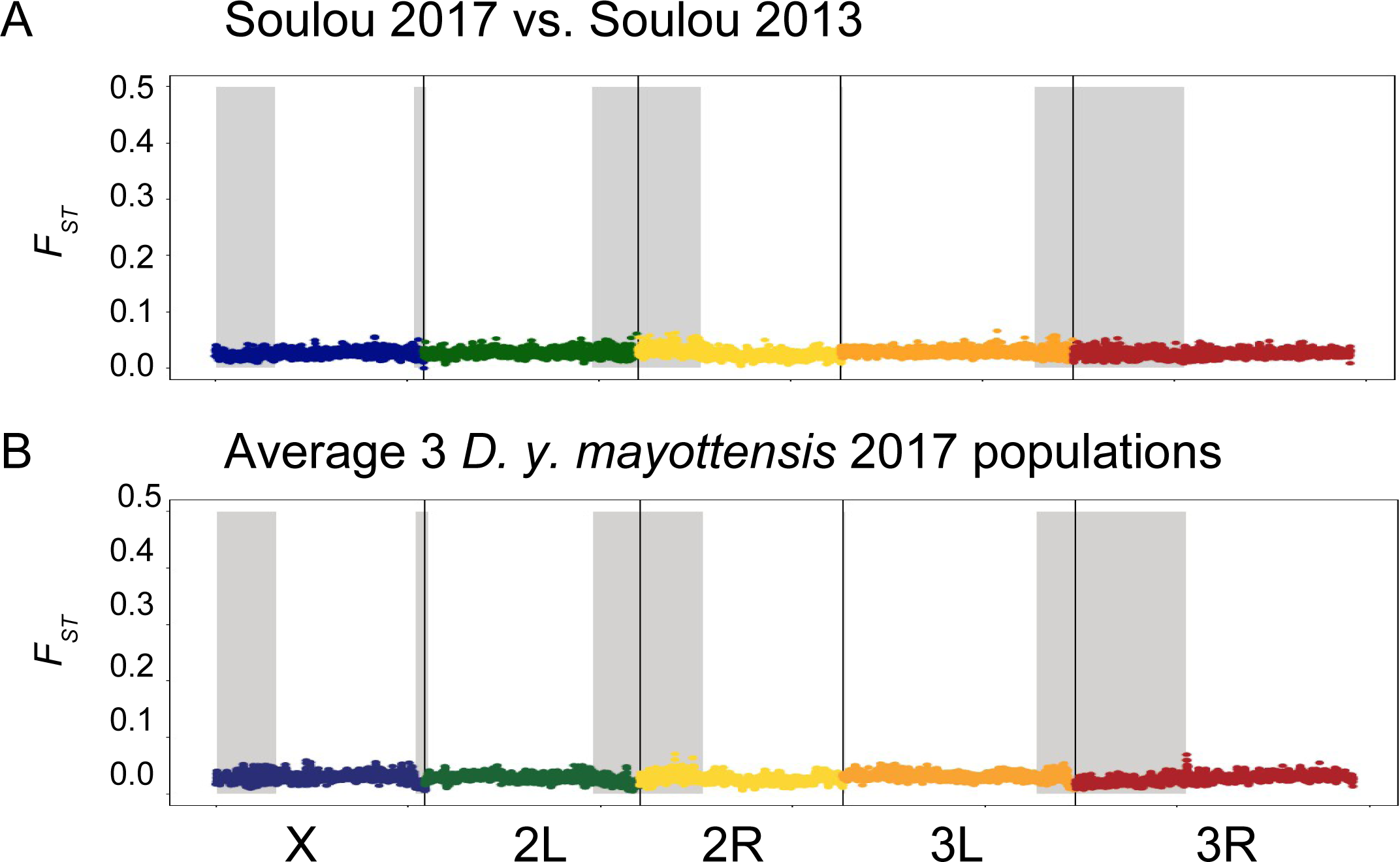
Low temporal and spatial genetic differentiation among *D. y. mayottensis* populations on the island of Mayotte. A) Pairwise *F_ST_* values for 10-kb windows between Pool-Seq data from Soulou collected in January 2013 and April 2017. B) Average pairwise *F_ST_* values for 10-kb windows between Pool-Seq data from the three populations collected in Mayotte in April 2017. Gray areas reflect subtelomeric and subcentromeric regions with low recombination rate (Yassin et al. 2016; Pettie et al. 2022). Each chromosomal arm according to the v.1.05 reference genome is represented by different color.

### Differentiation between the two *D. yakuba* subspecies differed among chromosomal arms

To quantify the level of genetic differentiation between Mayotte and mainland populations, we estimated *F_ST_* in 10-kb windows across the genome. Because of the lack of genetic structure among the three populations of *D. y. mayottensis*, we pooled either the Ind-Seq or the Pool-Seq sequences of the three populations in a single population, referring to them as Pool of Individuals (PoI) and Pool of Pools (PoP), respectively (Supplementary Table S2). Results did not greatly differ between the two methods, and we will show here analyses on the PoP data due to its larger sampling size and consequently reduced ascertainment bias. Surprisingly, the average genome-wide differentiation between the subspecies differed whether we used CY and NY, being higher for the former case (0.0731 ± 0.0003 and 0.0638 ± 0.0001, respectively; Student’s *t*-test *P* < 2.2 x 10^-16^). However, differentiation greatly depended on chromosomal arms (Figure 3A). Whereas for most chromosomal arms, differentiation between the two mainland populations CY and NY were significantly lower than between each from *D. y. mayottensis*, only for chromosomal arm 2R differentiation between CY and the island was the highest (Figure 3A). Differentiation among the three *D. y. mayottensis* populations did not show any chromosomal effect and for all chromosomal arms was significantly lower than differentiation between the two subspecies or within *D. y. yakuba* (Figure 3A; Supplementary Table S1).

**Figure 3 –.**
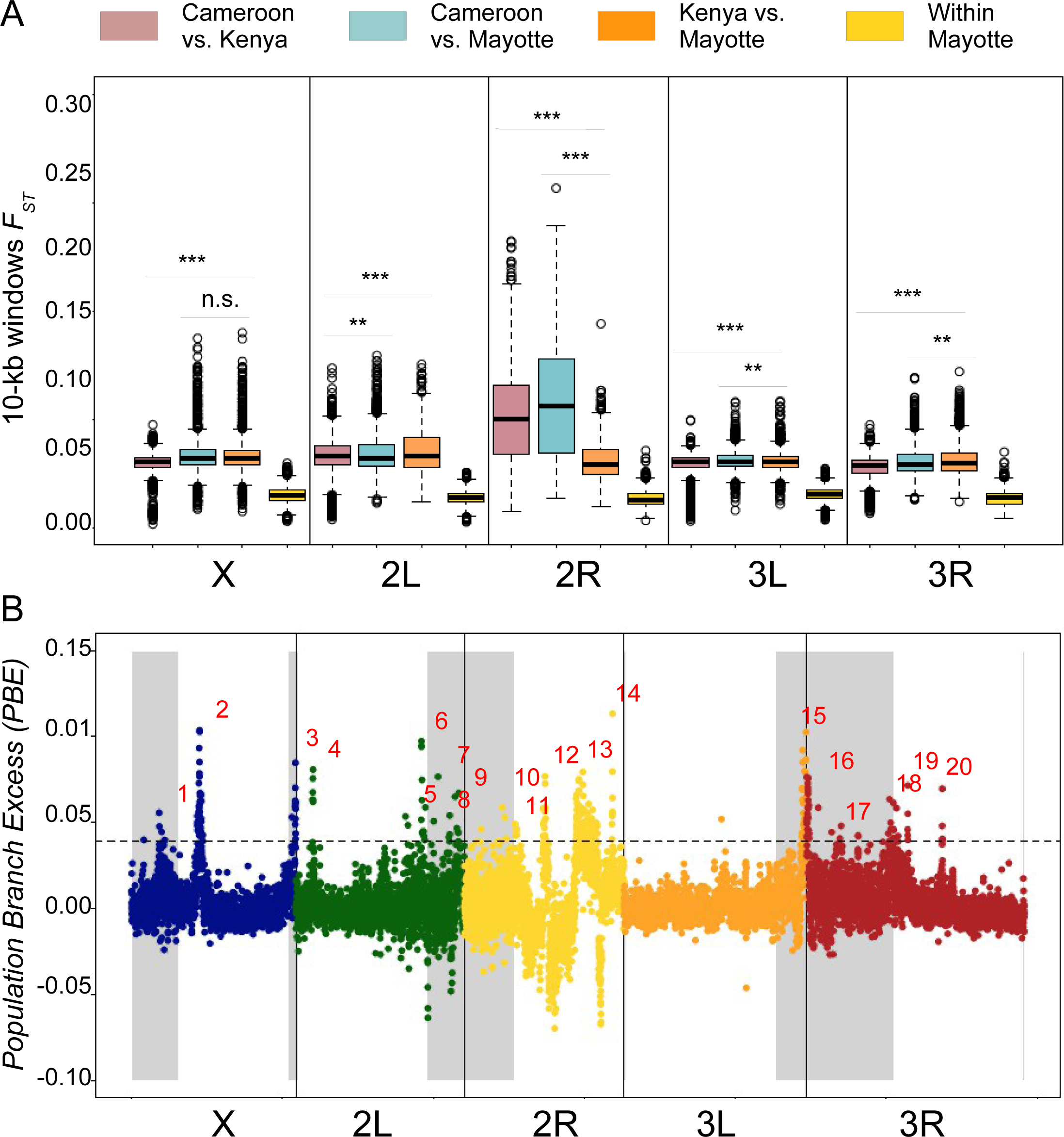
High genetic differentiation among the two *D. yakuba* subspecies. A) Boxplot of pairwise *F_ST_* values for 10-kb windows between the two mainland *D. y. yakuba* populations (pale red), between each and a pool-of-pooled flies from *D. y. mayottensis* (pale blue and orange), and between the three Pool-Seq populations of *D. y. mayottensis* (yellow). Significance estimated from Student’s *t* pairwise-tests. B) Population Branch Excess (*PBE*) identification of the genomic islands of divergence (GIDs), enumerated in red, distinguishing *D. y. mayottensis* from the two *D. y. yakuba* populations. Color code as in Figure 2.

### Twenty Genomic Islands of Divergence (GIDs) differentiate *D. y. mayottensis*

The *PBE* analysis showed 20 GIDs falling above the 97.5% quantiles in *D. y. mayottensis* (Figure 3B; Table 1). As in Yassin et al. (2016), we used the number of polymorphic sites found in 10-kb windows to delimit low recombining regions that are close to chromosomal arms extremities at the telomeres and centromeres (shaded areas in Figure 3B). These regions strongly concorded with fine-scale recombination rate variation in *D. yakuba* (Pettie et al. 2022). Half of the identified GIDs fell within these regions. Of the remaining 10 GIDs falling in high recombining regions, seven were on chromosome 2.

**Table 1 –.** Position, structural features and candidate genes or families of the 20 genomic islands of divergence (GIDs) distinguishing noni-specialist *Drosophila yakuba mayottensis* from generalist mainland species. *P* levels from DAVID gene ontology enrichment analyses for each GID.

| GI<br>D | Coordinates (Mb) | Architecture | Candidate genes |
| --- | --- | --- | --- |
| 1 | X:3.63-3.98 | Subtelomeric region |  |
| 2 | X:8.68-9.23 | Subspecies-specific *Xtb*<br>inversion | *Flo2*; Glucose-methanol-choline<br>oxidoreductase (ecdysteroid<br>metabolic process, $P < 1 \times 10^{-20}$) |
| 3 | X:21.22-21.77 | Subcentromeric region |  |
| 4 | 2L:2.21-2.27 | | Serine-type endopeptidase activity<br>($P < 3 \times 10^{-5}$) |
| 5 | 2L:14.51-14.83 | Subspecies-specific *2Lo*<br>inversion | *Adh* |
| 6 | 2L:16.59-17.15 | Subspecies-specific *2Lo*<br>inversion | *pdm3*; maltose alpha-glucosidase<br>activity (sugar metabolism and<br>transport, $P < 5 \times 10^{-13}$),<br>cytochrome P450 ($P < 0.10$) |
| 7 | 2L:18.24-18.83 | Subspecies-specific *2Lo*<br>inversion; subcentromeric<br>region |  |
| 8 | 2L:20.37-20.23 | Subspecies-specific *2Lo*<br>inversion; subcentromeric<br>region | *EcR* |
| 9 | 2L:21.08-21.57 | Close to subspecies-specific *2Lo* inversion;<br>subcentromeric region |  |
| 10 | 2R:5.01-5.57 | Subcentromeric region | *Catsup*; UDP-glycosyltransferases<br>(metabolism of xenobiotics by<br>cytochrome P450, $P < 0.01$) |
| 11 | 2R:6.85-6.93 | Close to species-common<br>*2Rj* inversion |  |
| 12 | 2R:10.42-10.77 | Species-common *2Rj*<br>inversion; subspecies-specific *2Rk* inversion | Cytochrome P450 ($P < 1 \times 10^{-5}$) |
| 13 | 2R:14.79-17.61 | Species-common *2Rj*<br>inversion; subspecies-specific *2Rk* inversion | LITAF ($P < 3 \times 10^{-14}$); glutathione<br>S-transferase ($P < 0.05$);<br>destabilase ($P < 0.05$); malic<br>enzyme ($P < 0.10$) |
| 14 | 2R:19.54-19.63 | Close to species-common<br>*2Rj* inversion |  |
| 15 | 3L:23.56-24.20 | Subspecies-specific *3Lk*<br>inversion; subcentromeric<br>region |  |
| 16 | 3R:0.00-0.33 | Subspecies-specific *3Ri* inversion; subcentromeric region |  |
| 17 | 3R:4.39-4.57 | Subcentromeric region |  |
| 18 | 3R:10.82-11.92 | Subcentromeric region | UDP-glucuronosyl/UDP-glucosyltransferase ($P < 5 \times 10^{-8}$) |
| 19 | 3R:13.38-13.42 |  | *Ace*, *Osi22* |
| 20 | 3R:17.96-17.98 | | *Osi2*, *Osi3*, *Osi4*, *Osi5* (Proteins of unknown function DUF1676, $P < 1 \times 10^{-8}$) |

Eight GIDs were enriched for specific gene families (Table 1). Gene families that were enriched for the whole list of genes from all GIDs included LITAF (LPS-induced TNF-activating factor) endosome-associated membrane proteins, glucose-methanol-choline (GMC) oxidoreductases, cytochrome P450, UDP-glycosyltransferases, and maltose-alpha-glucosidases at *P* < 0.05, and serine-type endopeptidases and glutathione-S-transferases at *P* < 0.10 (Supplementary Table S3). All these gene families have major roles in the metabolism of ecdysteroids, lipids, sugars, proteins and xenobiotics, as it would be expected in cases of major dietary shifts.

### GIDs in *D. y. mayottensis* predominantly overlap with chromosomal inversions

Our hybrid assembly of the *D. y. mayottensis* genome yielded 563 fragments with an N50 of 6.9Mb, and a BUSCOv5 score of 99.5%. Comparing the assembly to the v.1.05 reference genome assembly v.1.05 of *D. yakuba* identified 9 large structural variations with the characteristic V-shape of inversion breakpoints (Figure 4A). The first breakpoint was found on the X chromosome and indicated a ∼1 Mb-long inversion between X:8,648,348..9,652,060. Lemeunier and Ashburner (1976) did not detect any inversion on chromosome X in *D. y. yakuba*. Following their nomenclature, we call this new subspecies-specific inversion *Xto*, and note that it overlaps with GID2. We also noted a collinear connection between X: 13,477,771..17,070,823 on a single scaffold (grey) that could have resulted from double inversions, but do direct evidence for breakpoints was observed at this region (Figure 4A).

**Figure 4 –.**
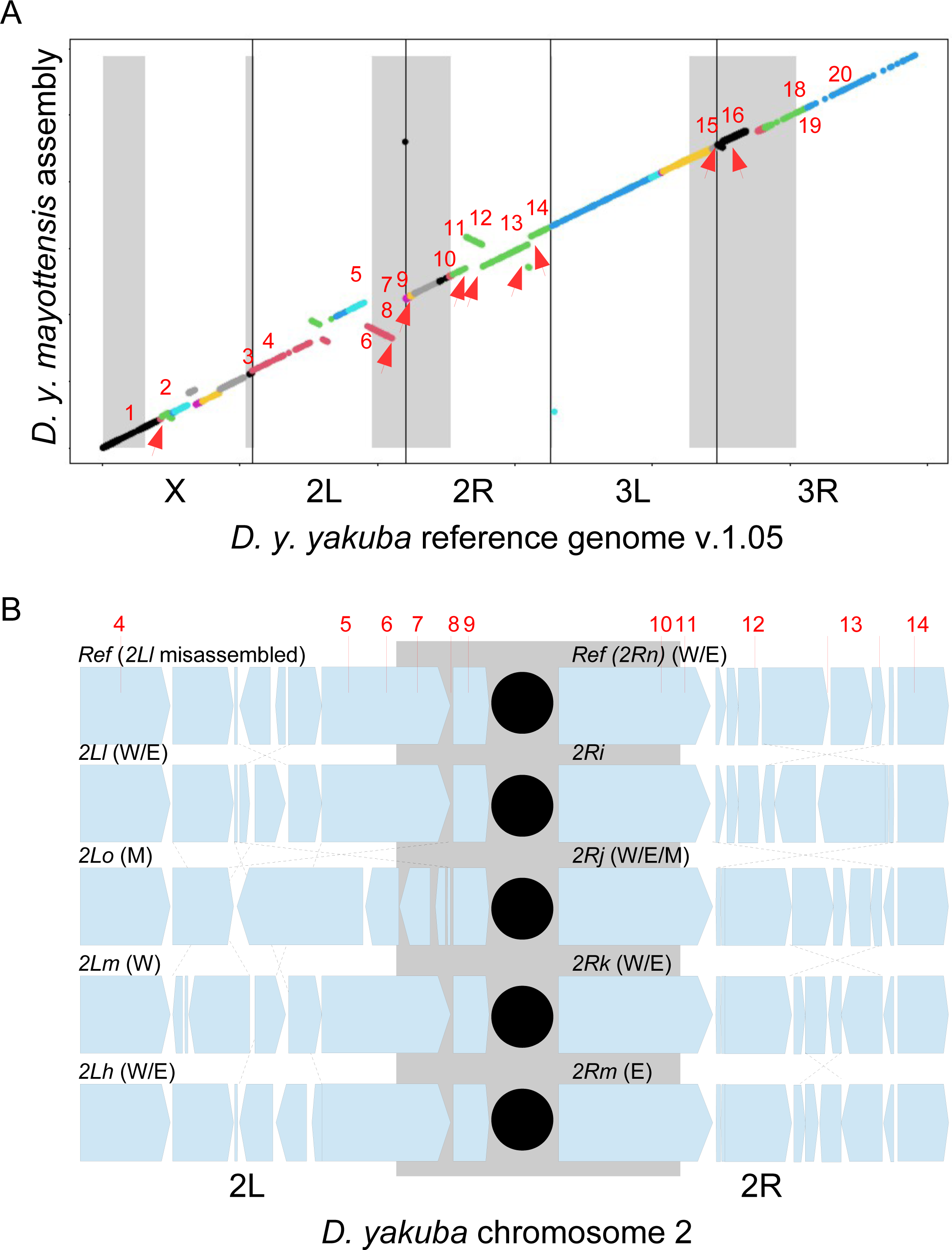
Genomic islands of divergence (GIDs) predominantly overlap with major chromosomal inversions. A) Dot plot comparing *D. y. mayottensis* hybrid assembly with the *D. yakuba* reference genome v1.05. Each *D. y. mayottensis* scaffold is given in different color with the position of the GIDs indicated by their number. Shaded areas indicate subtelomeric and subcentromeric regions with low recombination rate. Red arrows indicate inversion breakpoints. B) Schematic presentation of major inversions on chromosomal arms *2L* (left) and *2R* (right). Inversion names, breakpoints and geographical distribution into West Africa (W), East Africa (E) and Mayotte (M) are given following Lemeunier and Ashburner (1976) and this study.

The second breakpoint was found on chromosomal arm 2L and corresponded to a large, ∼12 Mb-long inversion between 2L: 8423810..20352247. This large inversion encompassed the misassembled portion of this chromosomal arm in the reference genome between 8,586,415 and 11,101,068 (Figure 4A,B; cf. Supplementary Figure S1 for alignment with a correct assembly). Lemeunier and Ashburner (1976) found four polymorphic inversions in mainland *D. y. yakuba*, namely *2Lh*, *2Ll*, *2Lm* and *2Ln*, with *2Lh* being ancestral to *2Ll*, and *2Lm* and *2Ln* independently derived from *2Ll*. We found the large inversion in *D. y. mayottensis* to be independently derived from *2Ll* too, and called it *2Lo* (Figure 4B). Remarkably, the proximal end of this inversion falls within the large subcentromeric region of 2L. Compared to the two most common inversions of the mainland, *i.e. 2Lh* and *2Ll* (Lemeunier and Ashburner 1976), four GIDs (5 to 8) fall in an inverted order in *2Lo* and one was close to its proximal, subcentromeric breakpoint (GID9) (Table 1).

The third breakpoint fell in the subcentromeric region of chromosomal arm 2R at 2R:182,816..436,892 but did not overlap with a major GID. However, the fourth to the seventh breakpoints all fell within the high recombining region of 2R. Their cytogenetic positions corresponded to the breakpoints separating chromosomal inversions *2Rn* and *2Rj* (Lemeunier and Ashburner 1976). The reference genome has the *2Rn* order indicating that *2Rj*, the most common inversion in *D. yakuba* (Lemeunier and Ashburner 1976), is fixed in *D. y. mayottensis*. Both inversions independently derived from a hypothetical *2Ri* order that has never been observed in natural populations (Figure 4B; Lemeunier and Ashburner 1976). Two GIDs, 11 and 14, fell in the flanking region separating *2Rj* from *2Ri*, whereas GID12 and 13 overlapped with the breakpoints separating *2Rn* from *2Ri*. Remarkably, the large GID13 is inverted in inversion *2Rk*, another common 2R inversion in *D. y. yakuba* that was not detected in *D. y. mayottensis*.

The eighth and ninth breakpoints were found in the subcentromeric regions of chromosomal arms 3L and 3R, respectively. Each corresponded to ∼0.2 and 0.8 Mb-long inversions and overlapped with GIDs 15 and 16, respectively. These small subcentromeric inversions were not detected by Lemeunier and Ashburner (1976) and we denoted them *3Lk* and *3Ri*, respectively.

## Discussion

By combining field observation with population and structural genomics, we were able to draw a clearer picture of the mechanisms underlying the evolution of genomic islands of divergence (GIDs) in the noni-specialist subspecies *Drosophila yakuba mayottensis*. First, we showed that *D. y. mayottensis* GIDs were subspecies-specific. There was no different pattern of divergence whether *F_ST_* was inferred by comparing mainland populations to the Soulou population collected in January 2013 or April 2017, or to any other noni-associated *D. y. mayottensis* population collected in 2017 across the island. Second, we found that the two subspecies had distinct inversions. Of the 20 GIDs distinguishing *D. y. mayottensis* from two mainland populations, 12 fell close to or within subspecies-specific chromosomal inversions, corresponding to ∼72% of the outlier windows in the *PBE* analysis.

Chromosomal inversions can play an important role in promoting local adaptation, either through reducing gene flow at selected genomic regions or through bringing to a new habitat an advantageous allelic combination segregating at the ancestral range (Faria et al. 2019; Mérot et al. 2020; Schaal et al. 2022). The two scenarios could have contributed to *D. y. mayottensis* specialization on noni in Mayotte. For example, GIDs falling in the subspecies-specific inversions *Xtb* and *2Lo* include several ecologically-relevant candidates that are involved in the metabolism of ecdysone, lipids and carbohydrates (GMC oxidoreductases (Iida et al. 2007; Glaser-Schmitt and Parsch 2018); maltases (Inomata et al. 2019)), ethanol tolerance (*Adh* (Siddiq and Thornton 2019)), and hexanoic acid sensing (*pdm3* (Arguello et al. 2021)). The *2Rj* inversion that is fixed in *D. y. mayottensis* is, on the other hand, widespread in *D. y. yakuba*, and it is ancestral to three 2R inversions that are specific to the mainland subspecies (Lemeunier and Ashburner 1976). GIDs near the breakpoints of this ancestral inversion contain important clusters of detoxification genes of cytochrome P450 oxidases and glutathione-S-transferases, which are involved in basal resistance to noni toxins in *D. sechellia* and its close relatives (Peyser et al. 2017; Lanno and Coolon 2019). Alternatively, both derived and shared inversions might have been fixed by drift, although demographic models do not suggest major bottlenecks during *D. y. mayottensis* colonization of Mayotte (Yassin et al. 2016). Testing the potential role of derived and shared inversions would require experimental isolation of each karyotype and comparison of the effect of each on different noni use traits, but the large breadth of these inversions would complicate fine-scale QTL mapping or genome-wide association studies (GWAS) in *D. yakuba*.

High levels of differentiation in collinear, high recombining regions constitute strong signals of hard selective sweeps. Only 3 GIDs (4, 19 and 20) satisfied these conditions. Most notably, GID20 consisted of syntenic members of the *Osiris* gene family. The role of those genes in conferring resistance against noni’s main toxin, octanoic acid, has been demonstrated through genetic mapping (Hungate et al. 2013) and transcriptomic analyses in *D. sechellia* (Lanno et al. 2017), as well as functional gene silencing in *D. melanogaster* (Andrade López et al. 2017; Lanno et al. 2019). The two other GIDs also involve some genes with a potential role in resistance to plant chemical defenses in other insects, such as *acetylcholine esterase* (*Ace*) or serine-type endopeptidases (Hoang et al. 2015; Alyokhin and Chen 2017; Lanno and Coolon 2019). Those genes therefore are best candidates for future functional genetics analyses. However, hard selective sweeps indicate a particular mode of selection wherein the selected alleles were absent or had low frequency at the onset of selection. Alternatively, incomplete selective sweeps wherein the selected alleles may have not reached fixation yet or soft selective sweeps on alleles segregating at intermediate frequencies in the ancestral range could also be important factors, although loci under these patterns of sweeps will hardly be detected by GID analyses. A signal of soft selective sweep on the odorant receptor *Or22a*, a gene that controls *D. sechellia* attraction to methyl hexanoate, the main noni volatile (Auer et al. 2020), was recently demonstrated in *D. y. mayottensis* (Ferreira et al. 2021). However, the efficiency of this approach will be limited to collinear, high recombining regions, which constitute a relatively small proportion of the *D. yakuba* genome.

In many cases, shared polymorphic inversions had promoted parallel adaptations in different geographical populations (Mérot et al. 2018; Lucek et al. 2019; Morales et al. 2019; Matschiner et al. 2022). This situation cannot explain the recurrent specialization on toxic noni fruits in distinct islands of the Indian Ocean between *D. yakuba* and *D. sechellia*. Despite their striking phenotypic convergence (Yassin et al. 2016; Ferreira et al. 2021), the divergence between the two species is deep and their genomic architectures are quite different. Nearly 35.3% of the genome in *D. yakuba* has low number of segregating sites extending from the telomeres and centromeres (Yassin et al. 2016; Pettie et al. 2022). In *D. simulans* and *D. mauritiana*, only 10.5% of the genome has low number of segregating sites (Brand et al. 2018). The three species of the *simulans* clade are homosequential and chromosomal inversions were very rarely reported in *D. simulans* (Chakraborty et al. 2021). As functional genetics analyses will continue to be undergone in both species, the degree of genic parallelism underlying the recurrent specialization on a toxic resource despite distinct genomic architectures will be clarified.

## Authors contributions

E.A.F. sequenced and assembled new genome, conducted population genomics and structural variation analyses, wrote the primary draft of the paper. C.C.M. sequenced individual lines. D.O. sequenced and assembled new genome. A.S. collected flies. S.R.P. taxonomically sorted flies. R.L.R. sequenced individual lines, obtained funds. A.Y. conceptualized the work, collected flies, taxonomically sorted flies, supervised population genomics and structural variation analyses, obtained funds, wrote the final version of the paper with inputs from all authors.

## Acknowledgments

The authors are grateful to Claire Golléty for help identifying noni plantation locations in Mayotte. Our late colleague, Jean R. David (1931-2021), who initially discovered *D. y. mayottensis* in 2013, participated to the 2017 expedition. This work was funded by the Agence Nationale de la Recherche grant no. ANR-18-CE2-0008, a Richard Lounsbery Foundation grant and an ATM bursary from the Muséum National d’Histoire Naturelle to AY, and a National Institutes of Health grant NIH NIGMS R35 GM133376 to RLR.

## Data availability statement

Scripts used in this study are publicly available on GitHub: https://github.com/AmirYassinLab/Population_genomics_scripts. Genome sequences generated for this study are available on NCBI BioProject: PRJNA972991, PRJNA973002 and PRJNA973271.

