## Supplementary figures and images for "Genomic islands of divergence between *Drosophila yakuba* subspecies predominantly overlap with chromosomal inversions"

### Supplementary Figure S1

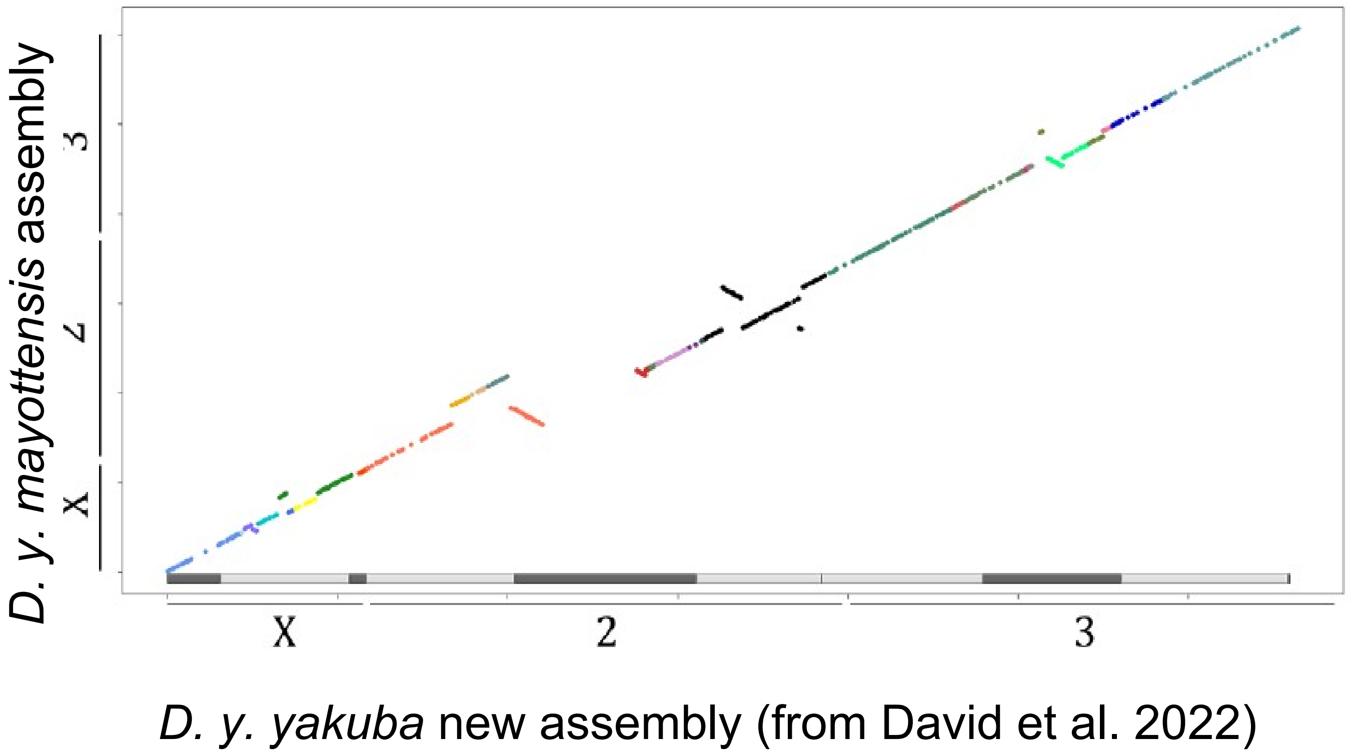
